# GOThresher: a program to remove annotation biases from protein function annotation datasets

**DOI:** 10.1101/2022.11.30.506803

**Authors:** Parnal Joshi, Sagnik Banerjee, Xiao Hu, Pranav M. Khade, Iddo Friedberg

## Abstract

**Motivation:** Advances in sequencing technologies have led to a surge in genomic data, although the functions of many gene products coded by these genes remain unknown. While in-depth, targeted experiments that determine the functions of these gene products are crucial and routinely performed, they fail to keep up with the inflow of novel genomic data. In an attempt to address this gap, high-throughput experiments are being conducted in which a large number of genes are investigated in a single study. The annotations generated as a result of these experiments are generally biased towards a small subset of less informative Gene Ontology (GO) terms. Identifying and removing biases from protein function annotation databases is important since biases impact our understanding of protein function by providing a poor picture of the annotation landscape. Additionally, as machine learning methods for predicting protein function are becoming increasingly prevalent, it is essential that they are trained on unbiased datasets. Therefore, it is not only crucial to be aware of biases, but also to judiciously remove them from annotation datasets.

**Results:** We introduce GOThresher, a Python tool that identifies and removes biases in function annotations from protein function annotation databases.

**Implementation and Availability:** GOThresher is written in Python and released via PyPI https://pypi.org/project/gothresher/ and on the Bioconda Anaconda channel https://anaconda.org/bioconda/gothresher. The source code is hosted on GitHub https://github.com/FriedbergLab/GOThresher and distributed under the GPL 3.0 license.

**Contact:** { | }

## 1 Introduction

The accurate annotation of the biological functions of proteins and other gene products is an open problem in life science. In the age of high throughput sequencing and multi-omics, accurate annotations are important: errors in initial annotations may propagate through different databases, leading to entrenched misannotations that are very difficult to root out (Schnoes *et al*., 2009). Furthermore, protein function prediction algorithms rely on functional annotation data as the standard-of-truth for training. It is commonly accepted that annotations using direct experimental evidence are of the highest quality. Biocurators typically assign function to gene products manually using the Gene Ontology (GO) (Ashburner *et al*., 2000) by examining experimental data from available literature, and assigning the appropriate GO terms to the relevant proteins (Camon *et al*., 2004). Given the standardized, dynamic and machine-readable format of Gene Ontology, it is currently the primary mode of annotating protein function.

In previous studies, we (Schnoes *et al*., 2013) and others (Clark and Radivojac, 2013; Bastian *et al*.,2015) have shown that not all experimental annotations are of equal quality. Specifically, experimental annotations from high-throughput studies appear to be less descriptive and contain less information than those from low throughput studies. An example of term with low information content is `catalytic activity (GO:0003824)` while a term with high information content would be *β*-alanyl-dopamine hydrolase activity (GO:0003832). The information content (IC) of a GO term is a quantitative measure that describes the specificity of that GO term within a certain context (Mazandu and Mulder, 2014). GOThresher calculates IC in the context of user provided set of annotations as the negative logarithm of the GO term frequency in the corpus. *IC_i_* = – log_2_(*P_i_*), where *P_i_* is the frequency of occurrence of any GO term *i* in a corpus as in (Lord *et al*., 2003; Pesquita *et al*., 2009). Annotations with low information content can be useful for some applications, such as genome-wide surveys, and they can provide insights into functioning of previously uncharacterized genes that lays groundwork for further experimental characterization (Attrill *et al*., 2019). However, such annotations do not accurately describe the function of a gene product. In response to the steady increase in number of high-throughput studies being published, the Gene Ontology Consortium introduced evidence codes that indicate annotations generated by high-throughput method as against other types of experiments which usually are small-scale and elucidate more detailed functionality about the gene products under investigation (Attrill *et al*., 2019). Moreover, computational methods trained on poor quality ground truth data generate annotations that are also non-specific, contributing further to the “shallow annotation problem” (Wang *et al*., 2007). Shallow annotations affect similarity measurements since gene pairs annotated exclusively with less informative terms like “protein binding” may demonstrate a non-meaningful functional similarity. However, this does not imply that these two gene products are similar in function, nor can these pairs be distinguished from other high-scoring gene pairs associated with more specific GO terms (Guzzi *et al*., 2012; Mistry and Pavlidis, 2008). This can also encourage algorithms trained on low information datasets will perform more shallow predictions (Clark and Radivojac, 2013). Other research has shown the how filtering large GO classes improves function prediction (Warwick Vesztrocy and Dessimoz, 2020; Törönen *et al*., 2018). In sum, a large number of low information content annotations that are mostly generated by large scale experimental studies strongly impact our understanding of protein function. To address the issue of bias arising from accumulation of less informative GO terms, we have developed GOThresher, which selectively removes annotations from Uniprot-GOA files based on GO evidence, annotation source, number of proteins annotated from a given source, date, or combinations of the above. GOThresher is useful to creators of tools for protein function prediction, for creating balanced training sets and selecting different levels of annotation specificity. GOThresher can be used standalone or in conjunction with protein function prediction benchmark creation tools such as GOBench (Dickson *et al*., 2022) or CAFA_Benchmark (https://github.com/CAFA-Challenge/CAFA_benchmark). It is also useful for data scientists interested in diagnosing protein function annotation databases.

## 2 Implementation

GOThresher is a command-line tool implemented in Python. We have released GOThresher as a package via PyPI and Conda enabling easy installation. GOThresher allows the user to obtain information about the quality of annotation datasets using Information Content (IC) calculation as described in (Lord *et al*., 2003) and Information Accretion as described in (Clark and Radivojac, 2013). The main functionality that GOThresher provides is to remedy biases from protein function annotation datasets using filtering criteria based on annotation information content, annotation source, and several proteins annotated by a certain source. GOThresher requires Gene Ontology in the Open Biomedical Ontologies (OBO) format and protein function annotation data in Gene Association File (GAF) format (Gene-Ontology-Consortium, 2012).

### Select functions of GOThresher

For a full list of features, see the documentation on the Github site. Using the -PLTHRESH argument, the user can provide a threshold for discarding annotations with an IC below it. Instead of providing a value, it is also possible to provide a percentage value *p* following the -PLTHRESHp argument, which will retain only those annotations whose IC is in the top *p*% of all the IC scores.

### Example usage

~~~
$ gothresher -i goa_yeast.gaf -a C P -e EXPEC IBA -PLTHRESH 5 -o results_dir
~~~

This command will read goa_yeast.gaf, select the Cellular Component and Biological Process GO terms which have been determined experimentally or generated computationally as “IBA” (Inferred from Biological Aspect of ancestor). Additionally, it will discard all annotations that have an IC ≤ 5 and create the output in the `results_dir` directory.

## 3 Results

Here we show some results illustrating the utility of GOThresher in filtering information from a UniProt GOA file. The results demonstrate the utility of GOThresher in balancing the databases.

**Input data** (see GitHub repository: https://github.com/FriedbergLab/GOThresher): goa_exampleYeast.gaf which is a truncated Uniprot-GOA file based on 2022-01-14 version of the yeast Uniprot-GOA file available in full from: https://ftp.ebi.ac.uk/pub/databases/GO/goa/old/YEAST/

### Example 1: Filter by Information Content (IC)

**Command:**

~~~
$ gothresher -PLTHRESH 10 -i goa_exampleYeast.gaf
~~~

**Explanation:** This command will read `goa_exampleYeast.gaf`, discard all the annotations that have an IC ≤10, and create an output file named `exampleYeast_MFO_BPO_CCO_PL_10.gaf` in the working directory.

#### Example of processing

Maximum IC of GO term in the input data before filtering: 15.5

Minimum IC of GO term in the input data before filtering: 5.2

IC threshold to discard GO terms: 10

**Table S1:**
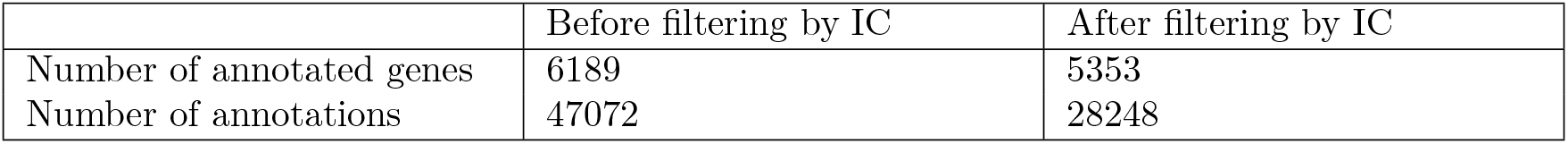
Difference between number of annotated genes and all annotations before and after filtering based on Philip Lord Information Content

|  | Before filtering by IC | After filtering by IC |
| --- | --- | --- |
| Number of annotated genes | 6189 | 5353 |
| Number of annotations | 47072 | 28248 |

**Figure S1:**
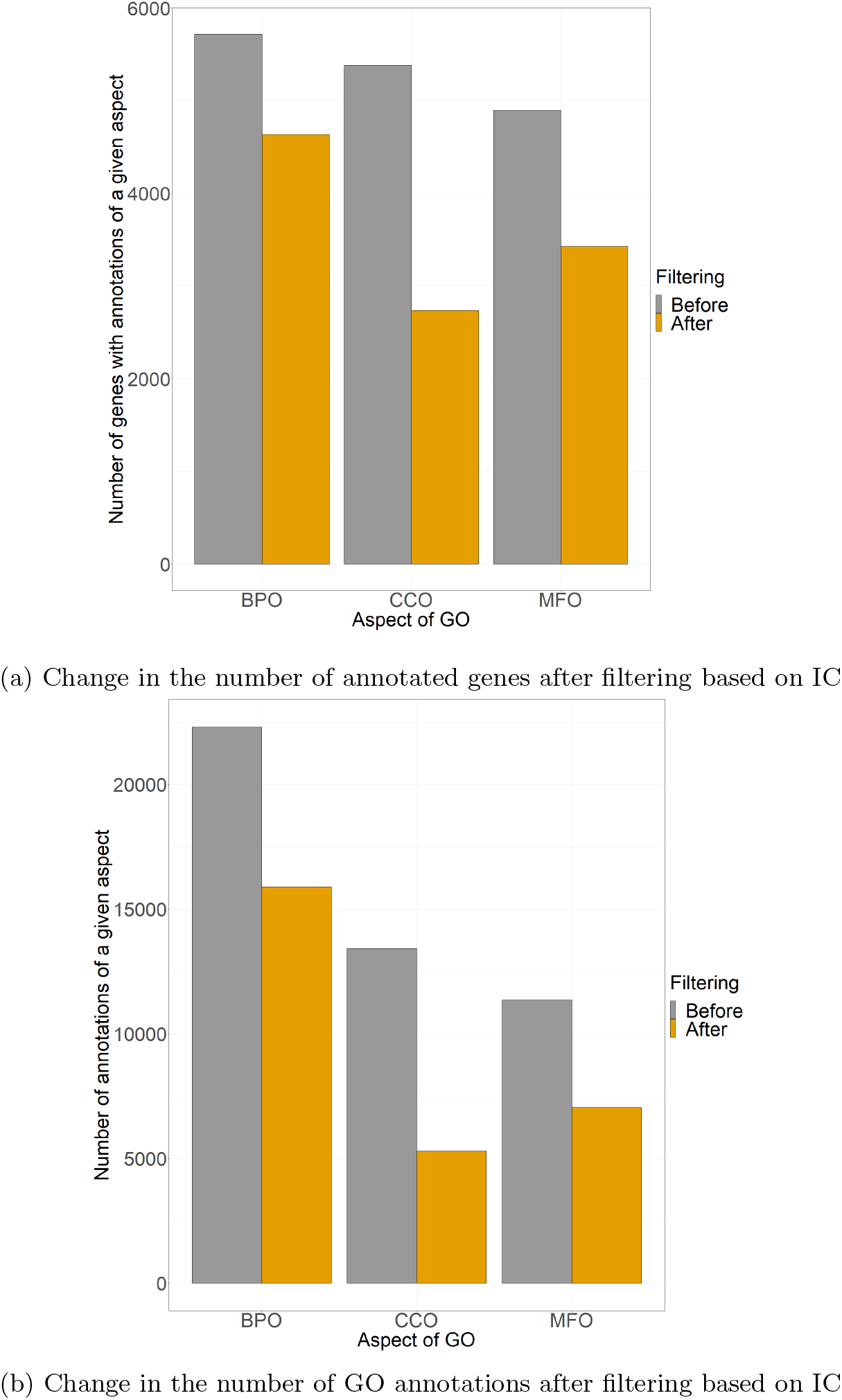
Difference between the (a) number of annotated genes and (b) number annotations separated by the three GO aspects Biological Process Ontology (BPO), Cellular Component Ontology (CCO), and Molecular Function Ontology (MFO) before and after filtering based on Philip Lord Information Content

### Example 2: Filter by evidence code – select all experimental annotations

**Command:**

~~~
$ gothresher -e EXPEC -i goa_exampleYeast.gaf
~~~

**Explanation:** This command reads `goa_exampleYeast.gaf`, selects annotations that have been generated experimentally, and creates an output file named exampleYeast_MFO_BPO_CCO_EXPEC.gaf in the working directory. Experimental evidence includes the following GO experimental evidence codes: [‘EXP’, ‘IDA’, ‘IPI’, ‘IMP’, ‘IGI’, ‘IEP’, ‘HTP’, ‘HDA’, ‘HMP’, ‘HGI’, ‘HEP’] (See http://geneontology.org/docs/guide-go-evidence-codes/ for an explanation of evidence codes)

**Table S2:** Difference between number of annotated genes and all annotations before and after filtering based on experimental evidence

|  | Before filtering by evidence | After filtering by evidence |
| --- | --- | --- |
| Number of annotated genes | 6189 | 5827 |
| Number of annotations | 47072 | 42930 |

**Figure S2:**
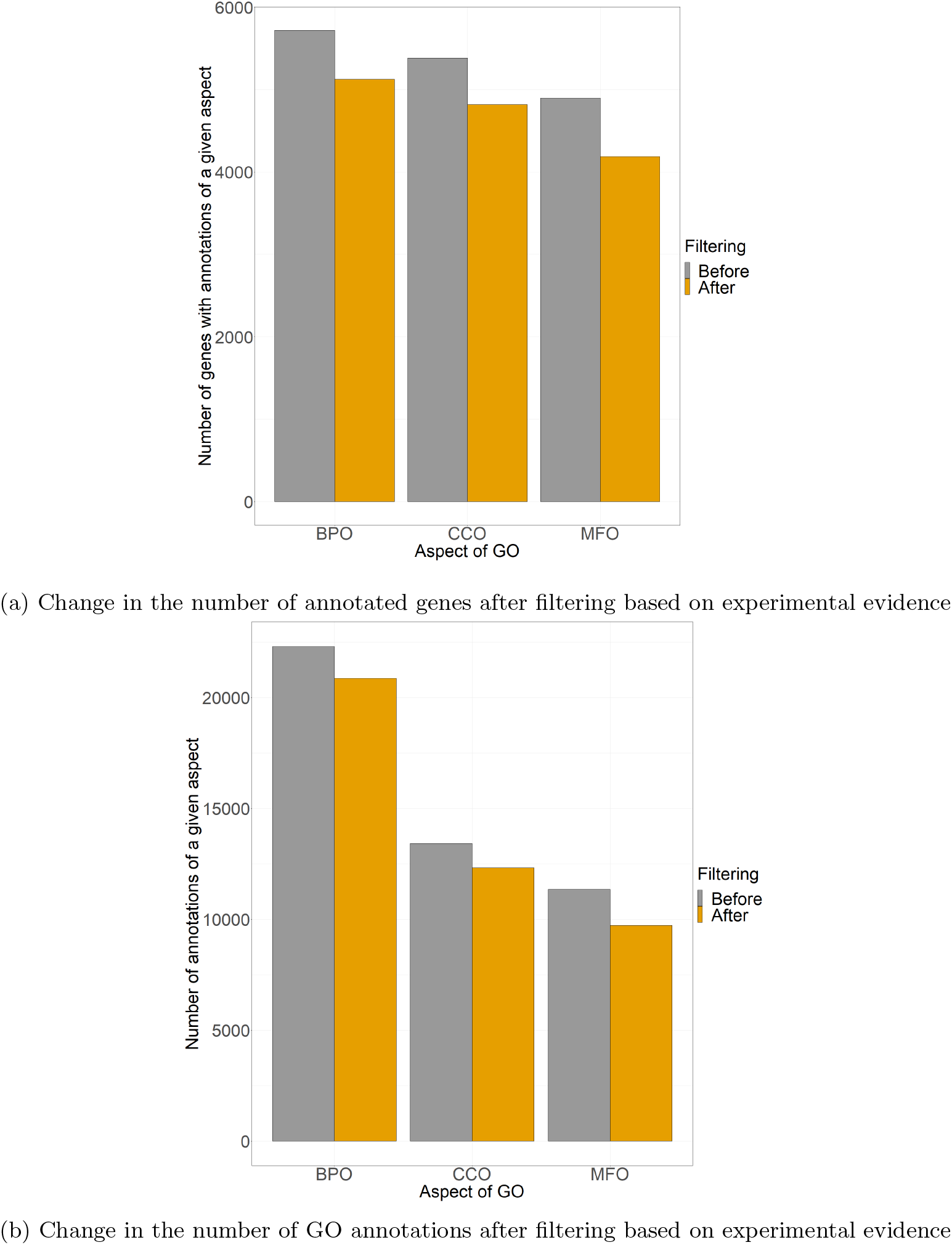
Difference between the (a) number of annotated genes and (b) number of annotations separated by the three GO aspects Biological Process Ontology (BPO), Cellular Component Ontology (CCO), and Molecular Function Ontology (MFO) before and after filtering using experimental evidence

### Example 3: Filter by aspect of GO – select Molecular Function Ontology (MFO) GO terms not generated experimentally

**Command:**

`$ gothresher -einv EXPEC -a F -i goa_exampleYeast.gaf`

**Explanation:** This command reads `goa_exampleYeast.gaf`, discards annotations that are associated with all experimental evidence codes, then selects Molecular Function GO terms, and creates an output file named `exampleYeast_MFO.gaf` in the working directory.

**Table S3:**
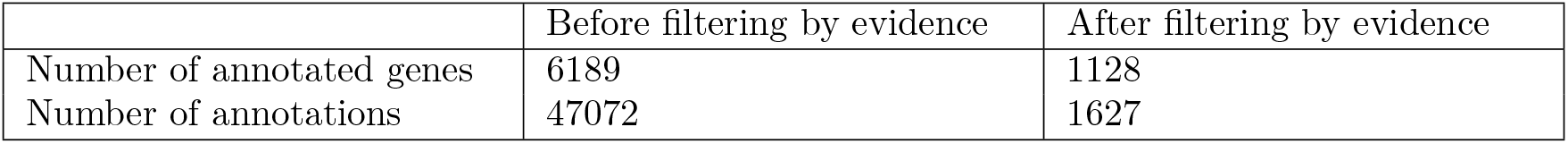
Difference between number of annotated genes and all annotations before and after filtering based on aspect of GO and evidence

|  | Before filtering by evidence | After filtering by evidence |
| --- | --- | --- |
| Number of annotated genes | 6189 | 1128 |
| Number of annotations | 47072 | 1627 |

**Figure S3:**
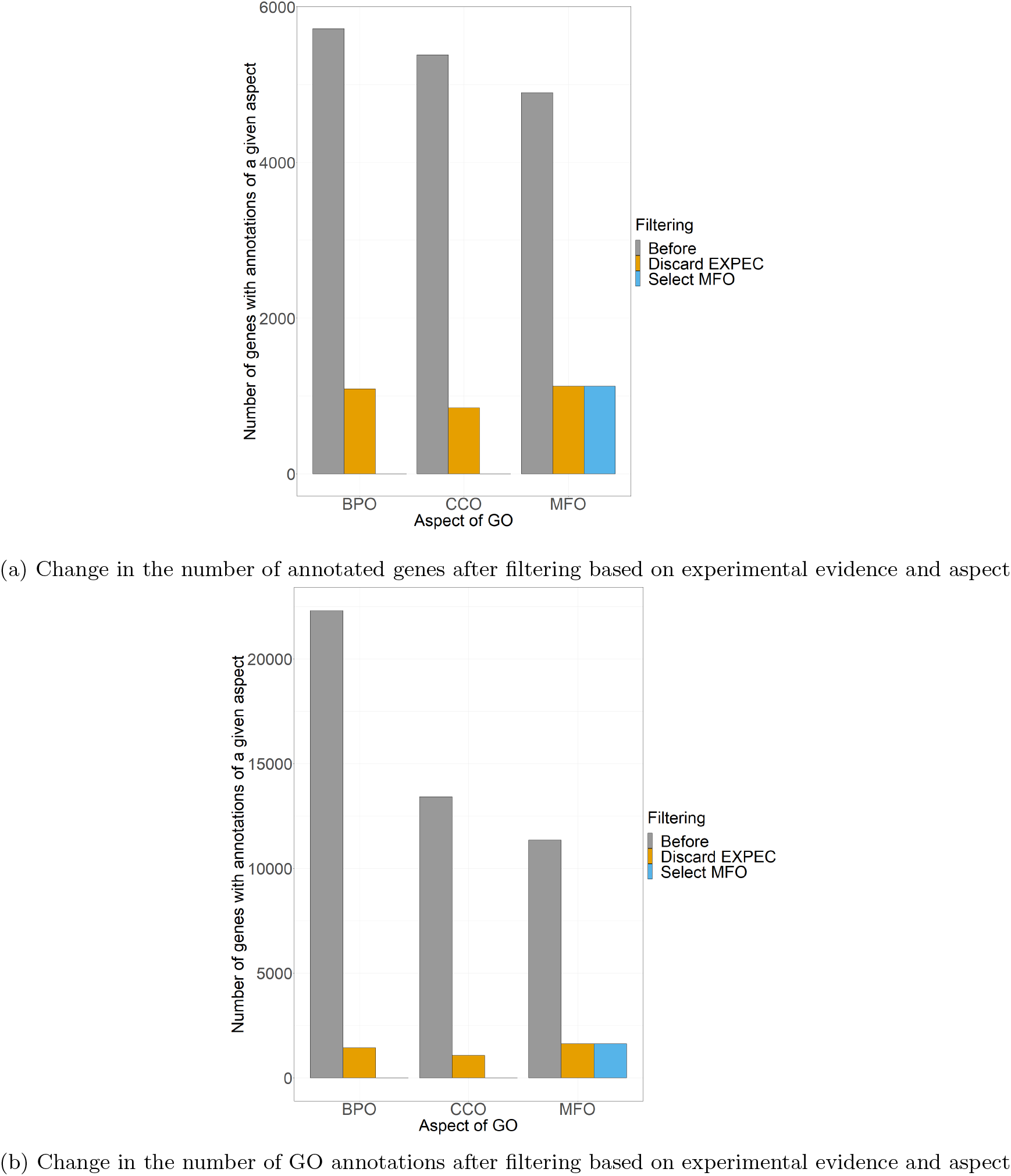
Difference between the (a) number of annotated genes and (b) number of annotations separated by the three GO aspects Biological Process Ontology (BPO), Cellular Component Ontology (CCO), and Molecular Function Ontology (MFO) before and after filtering based on the aspect of GO and evidence

